# Postpartum trend in blood pressure and renal function in women with severe preeclampsia and eclampsia: A prospective cohort-study at Mulago hospital, Kampala, Uganda

**DOI:** 10.1101/562934

**Authors:** Kasereka Muteke, Jolly Beyeza, Milton W Musaba, Julius Wandabwa, Paul Kiondo

**Affiliations:** Department of Obstetrics and gynecology, College of Health Sciences, Makerere University, P.O BOX 7072, Kampala, Uganda, East Africa.; Department of Obstetrics and Gynecology, Mulago National Referral Hospital, P.O Box 7051, Kampala, Uganda, East Africa; Department of Obstetrics and Gynecology, Faculty of Health Sciences, P.O Box 1460, Mbale, Uganda

**Keywords:** Preeclampsia, Hypertension, Proteinuria, Renal dysfunction, Resolution, Postpartum

## Abstract

**Background:** Preeclampsia/Eclampsia is a multisystem disorder of pregnancy with kidney involvement. Our objective was to assess the postpartum trend in blood pressure, renal function and proteinuria and, to investigate their predictors in Ugandan women with severe preeclampsia and eclampsia.

**Methods:** This was a prospective cohort study that involved 97 women with severe preeclampsia and Eclampsia, conducted at Mulago National referral hospital from August 2017 to April 2018. The clinical and laboratory variables were collected from the women on day1, 7, 21 and day 42 after delivery. Kaplan-Meier Survival analysis, Cox-proportional Regression and Log-Rank tests were used to compare the baseline and the follow-up variables with changes in blood pressure, renal function and urine protein.

**Results:** Most women (93.8%) recovered from hypertension within 6 weeks of childbirth with the mean time to resolution of 2.49 weeks (95% CI: 2.13-2.82). About 81% of the women recovered their renal function and the mean time to recovery was 24.54 days (95% CI: 20.14-28.95). Proteinuria resolved in approximately 84% of the women and the mean time resolution of urine protein of 32.85 days (95% CI: 30.31-35.39). Having multiple pregnancy versus a singleton pregnancy was associated with persistence of hypertension six weeks after child birth (P-value = 0.013).

**Conclusion:** In this study, the blood pressure and renal function of most women with severe preeclampsia and eclampsia normalized within six weeks after childbirth. A special interdisciplinary follow up for patients with preeclampsia/eclampsia by an obstetrician and physician is needed in the postpartum period to reduce the maternal morbidity and mortality associated with this condition in our community.

## Introduction

Preeclampsia is a multisystem human specific pregnancy disorder characterized by new onset hypertension and proteinuria after 20 weeks of pregnancy(1). It affects 2-8% of all pregnancies worldwide and contributes significantly to maternal, fetal and neonatal morbidity and mortality (2). Preeclampsia with other hypertensive disorders in pregnancy contributed to 14% of maternal deaths worldwide (3). It is estimated to cause 8% of the severe maternal morbidity in Uganda is the leading cause of maternal deaths (4). Women with preeclampsia have an increased risk of renal, cerebrovascular and cardiovascular complications after delivery(5). In low resource settings, preeclampsia is an important cause acute kidney injury and contributes to one third of the cases seen in late pregnancy (6). Half of the women with acute kidney injury require dialysis and when dialysis is not available as is commonly the case in many low resource settings, acute kidney injury frequently leads to death of the women. Studies show that women recover their renal function after preeclampsia (7), however, other workers have revealed that women with preeclampsia are at a 5 to 12-fold increased risk of end-stage renal disease (8) and therefore require prolonged nephrological follow up. The development of renal disease after preeclampsia is not clearly understood. The renal injury may be due to extensive endothelial or podocyte injury (9) seen in women with preeclampsia. This leads to nephron loss and later development of renal disease.

Women with preeclampsia on the other hand are at increased risk of cardiovascular disease compared to normotensive women. They therefore require long time follow up regarding hypertension after delivery(10). The mechanisms of developing chronic hypertension is not clear. However, it may be due organ damage or preeclampsia may be a risk factor for later development chronic hypertension. (1). Studies in Uganda have shown that up to one third of the women with preeclampsia had persistent hypertension after the puerperium (11, 12). The predictors for persistent hypertension were participants age (11, 12), gestational age at delivery and parity of the mother (12).

It is therefore important that women with preeclampsia are followed after the puerperium if the blood pressure and renal dysfunction do not resolve. The purpose of this study therefore was to evaluate the postpartum trend in blood pressure, renal dysfunction and, proteinuria and to determine the factors associated with their resolution in women with severe preeclampsia and Eclampsia in Mulago hospital.

## Methods

### Study design

This was a prospective cohort study conducted from August 2017 to April 2018 in Uganda. This study was approved by the Mulago hospital ethics committee, the Makerere School of Medicine Research and Ethics committee and Uganda National Council for Science and Technology. Written informed consent was obtained from the participants.

### Setting

This study was conducted at Mulago hospital. Mulago hospital is a national referral hospital for Uganda and serves as the teaching hospital for Makerere University College of Health Sciences. Mulago Hospital delivers about 30,000 women per year and offers antenatal and postnatal services.

### Study population

The study population consisted of women with severe preeclampsia and eclampsia who delivered at Mulago hospital during the study period. Women with a known history of hypertension, diabetes mellitus and kidney disease were excluded from the study.

Preeclampsia was defined according to the classification by the Working group of National High blood pressure Education program (2000) and the American College of Obstetricians and Gynecologists (2013) (13). Under this classification, hypertension was defined as a systolic blood pressure of ≥ 140mmHg and/or diastolic blood pressure of ≥ 90mmHg on 2 occasions at least 4 hours or more apart. Proteinuria was defined as urine protein of ≥300mg/24h urine collection or protein/creatinine ratio of ≥ 0.3 or a dipstick reading of ≥2+. Preeclampsia was taken as hypertension with proteinuria after 20 weeks gestation.

A woman with preeclampsia was taken to have severe preeclampsia if she had BP of ≥160 mmHg systolic or ≥110 mmHg diastolic, severe headache or visual disturbances, thrombocytopenia of ≤100,000/µL, aspartate transaminase or alanine transaminase > 2times the upper limit with severe epigastric or upper quadrant pain, pulmonary edema or serum creatinine ≥1.1mg/dL. A woman with preeclampsia was taken to have eclampsia if she developed a convulsion that could not be attributed to any other cause (14).

### Sample size

We assumed that the persistence of hypertension would be 42.6% as was found in a study by Kaze et.al (15) and parity as a biggest risk factor for preeclampsia with an odds ratio of 3.71 as was found in a study in Mulago hospital (16). With these estimates a sample size of 97 participants would be sufficient with power of 80% at confidence level of 95% taking in account of the anticipated loss to follow up of 5%.

### Study procedures

The research assistants who were qualified midwives identified women with severe preeclampsia and eclampsia from the labour ward and the high dependence unit of the hospital. They approached the attendants of the women and gave them information about the study. The attendants were conducted through an informed consent procedure and gave a written informed consent. The women later gave informed consent when they improved. Information was obtained from the attendants and from the abstraction of the charts and later verified from the women when they improved. The eligible participants were recruited consecutively until the required sample size was achieved. The information from the women was collected using an interviewer-administered questionnaire, participants’ examination, and biochemical investigations. Urine was collected from the women for urine protein estimation and blood was drawn for serum creatinine measurement.

### Follow up

The women were followed for 6 weeks after delivery. The women were reviewed on day 1,7,21 and day 42 by the research team. During the review the women were asked about their health and a focused history and examination were done using case record form. The blood pressure was measured, blood was drawn from the women for measurement of serum creatinine and urine for estimation of urine protein. For renal function, a software package was used to estimate the glomerular filtration rate using current serum creatinine, patient race, gender and age of the participant (17). The MDRD calculator was used for determining and classifying the estimated glomerular filtration https://patient.info/doctor/estimated-glomerular-filtration-rate-gfr-calculator

### Outcomes

The primary outcome was time to resolution of the blood pressure. Secondary outcomes were time to recovery of renal function and disappearance of proteinuria. The blood pressure was considered normal when it was less than 140/90 mmHg without any antihypertensive medications for at least one week.

Serum creatinine level and estimated glomerular filtration were measured at every visit and were considered normal when the estimated glomerular filtration was ≥ 90 mL/min/1.73 m^2^ Urine protein was considered to have returned to normal if the measurement of spot urine by dipstick was less than 30mg/dl.

### Data analysis

The data were coded and double entered using the Epi-data Version 3.1 and analyzed with STATA version 12. Counts, means, median, percentage and cumulative percentage were used to report the results. Survival analysis with Kaplan-Meier was used to determine time to resolution of hypertension, renal dysfunction and urine protein. Cox- proportional regression and Log rank test were used to determine association of participant variables with time-to resolution of hypertension, renal dysfunction and urine protein. Association was considered statistically significant if it had a P-value of less than 0.05.

## Results

In this study 97 women with severe preeclampsia/Eclampsia were followed up for 6 weeks after childbirth. The time to resolution of hypertension, renal function and proteinuria, and the associated factors were determined.

All the 97 participants had hypertension, 20 were censored: 2 women died, 12 were lost to follow-up by the third visit, and 6 had persistent hypertension by the end of this study.

Forty seven participants had abnormal renal functions, 10 were censored: 9 women had persistent renal dysfunction by the end of the study and 1 woman was lost follow up.

Finally, 92 had proteinuria: 29 were censored because 15 women had persistent proteinuria by the end of the study and 14 women were lost to follow up (Fig 1).

**Figure 1:**
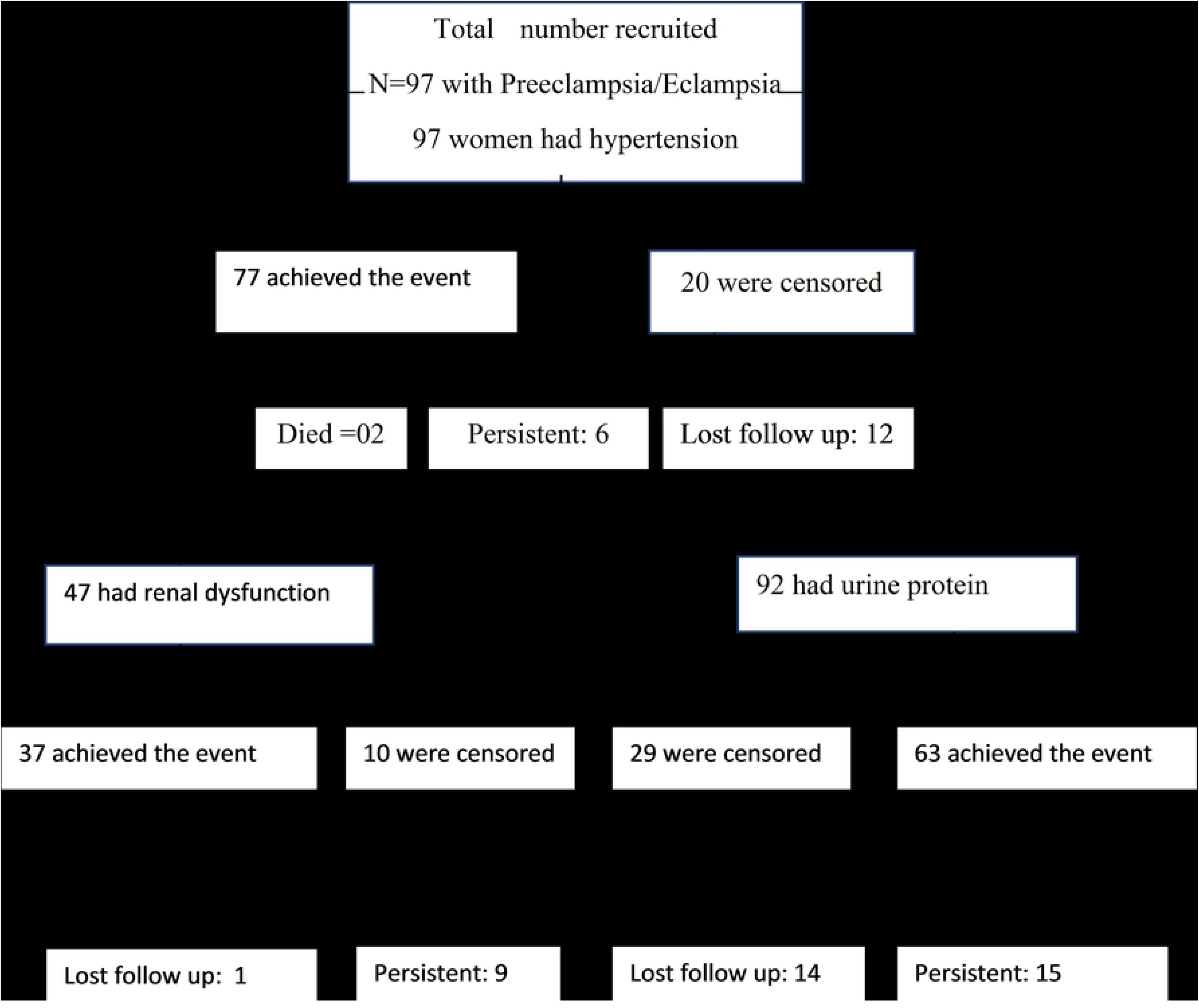
Flow chat of 97 participants with severe preeclampsia and eclampsia.

The mean age of the participants was 26.6±5.4 years, mean gestation age was 35.9±4.0 weeks and a modal parity was 2 with a range of 1-6.

The mean time to resolution of hypertension was 2.49 weeks (CI: 2.13-2.84). The blood pressure decreased over the 6 weeks period of follow-up. Only 6 women (6.2%) had persistent hypertension 6 weeks after delivery (Fig 2). The decrease of blood pressure was not affected by the time of onset of preeclampsia (p=0.426), mode of delivery (p=0.891) and the parity of the woman (p=0.139).

**Figure 2:**
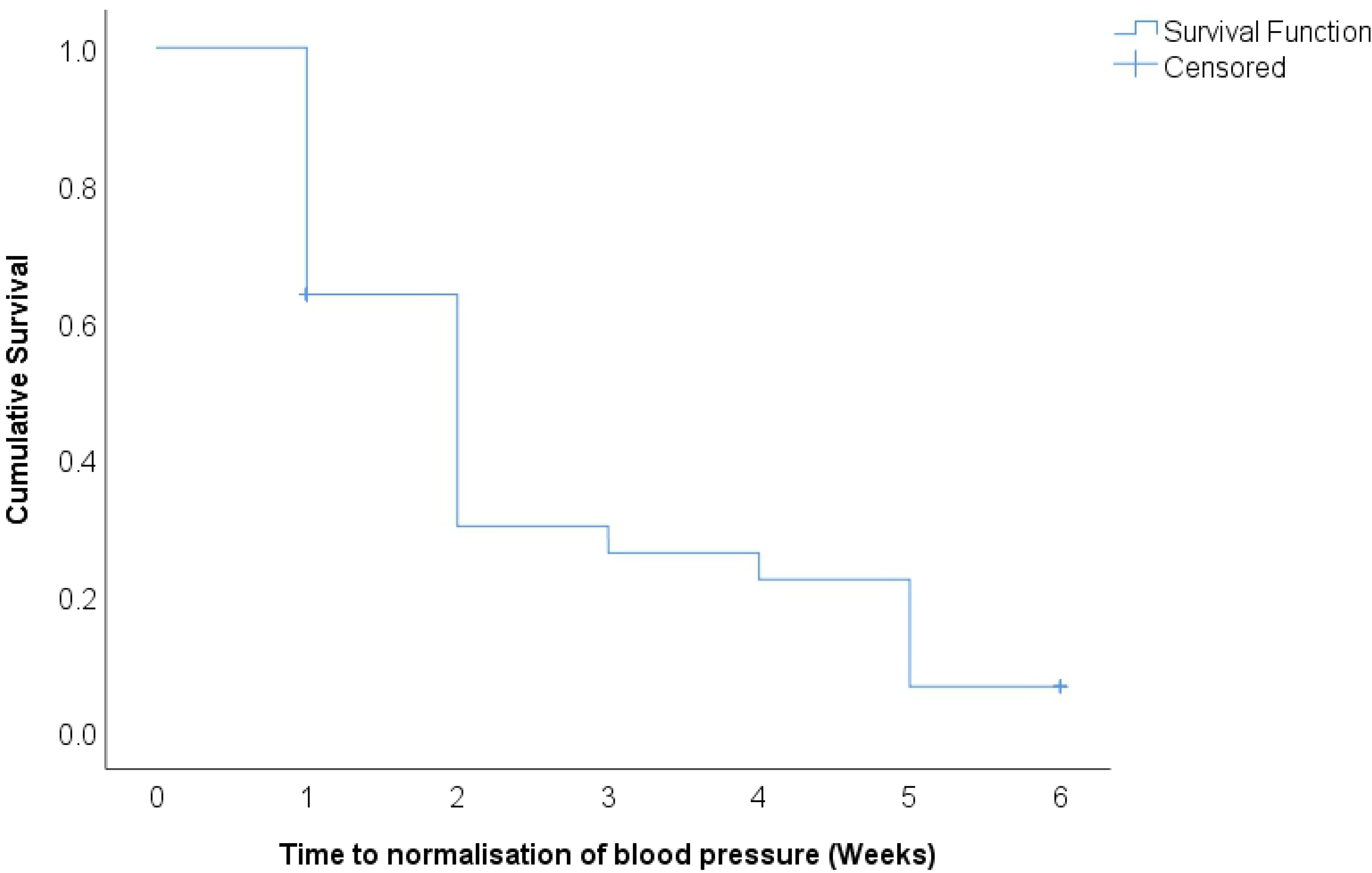
Kaplan-Meier curve of time to resolution of hypertension in women with preeclampsia and eclampsia.

After controlling for other variables, having multiple pregnancy versus singleton pregnancy was significantly associated with persistence of hypertension 6 weeks after delivery (p value =0.013) (Fig 3). Other factors like time of development of preeclampsia, parity and mode of delivery were not associated with persistence of hypertension 6 weeks after childbirth.

**Figure 3:**
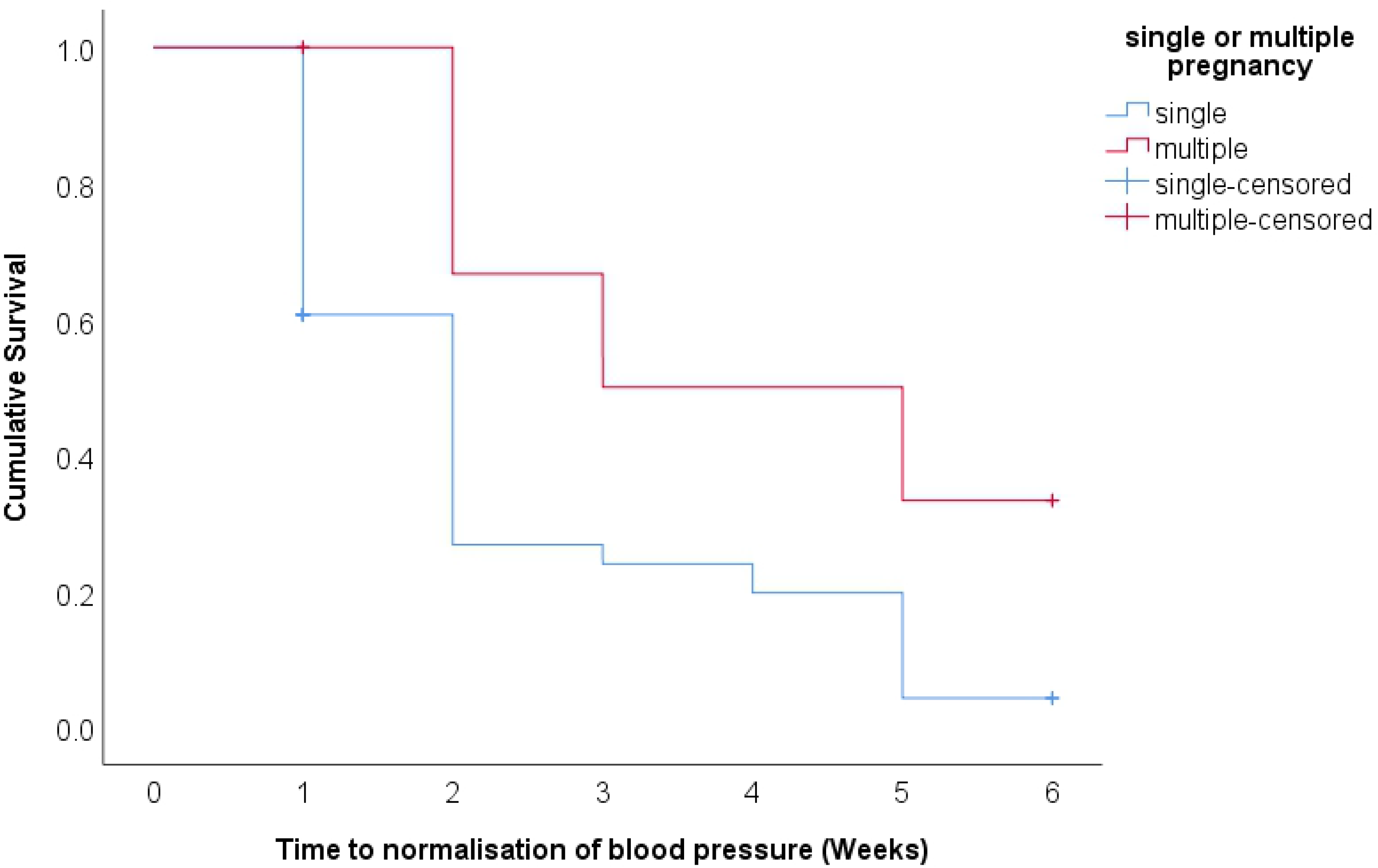
Kaplan-Meier curve for time to resolution of hypertension in women with single versus multiple pregnancy.

The mean time for the recovery of the renal function was 24.5 days (95% CI: 20.14-28.95). The renal function improved during the six weeks of follow up and only 9 (19.1%) women had persistent renal dysfunction at 6 weeks follow-up (Fig 4). There recovery of the renal function was not associated with mode of delivery of the patient (p=0.256), onset of preeclampsia (p=0.180), parity of the patent (p=0.709) and whether the mother delivered single or multiple pregnancy (p=0.147).

**Figure 4:**
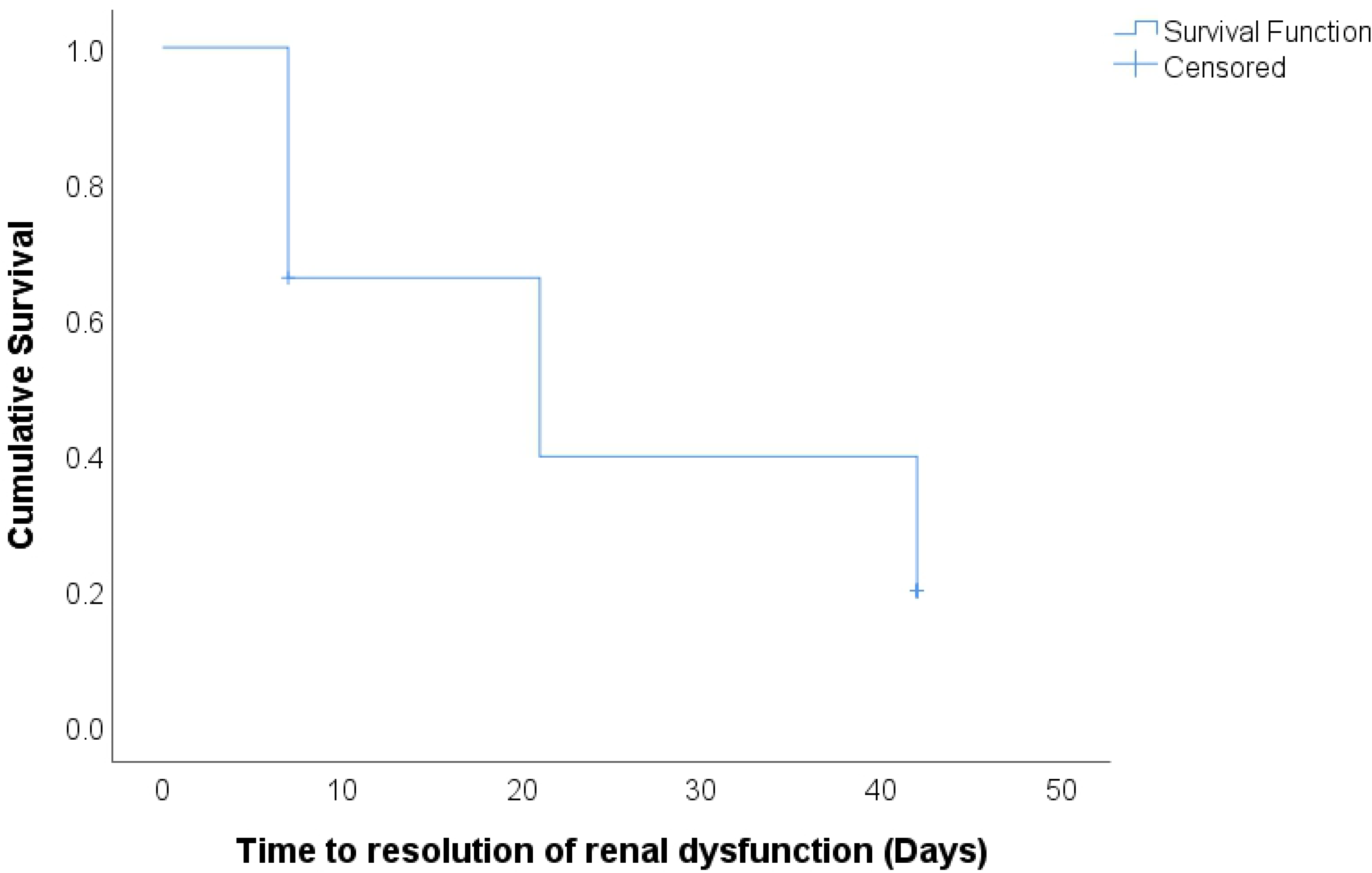
Kaplan-Meier curve of time to resolution of renal dysfunction in women with severe preeclampsia and eclampsia.

The mean time for the resolution of proteinuria was 32.9 days (95% CI: 30.3-35.4). Urine protein decreased over the six week follow-up and 15 (16.3%) women had persistent proteinuria by the end of the study (Fig 5). The resolution of proteinuria was not associated with mode of delivery (p=0.267), onset of preeclampsia (p=0.660), parity (p=0.135) and single versus multiple pregnancy (p=0.075).

**Figure 5:**
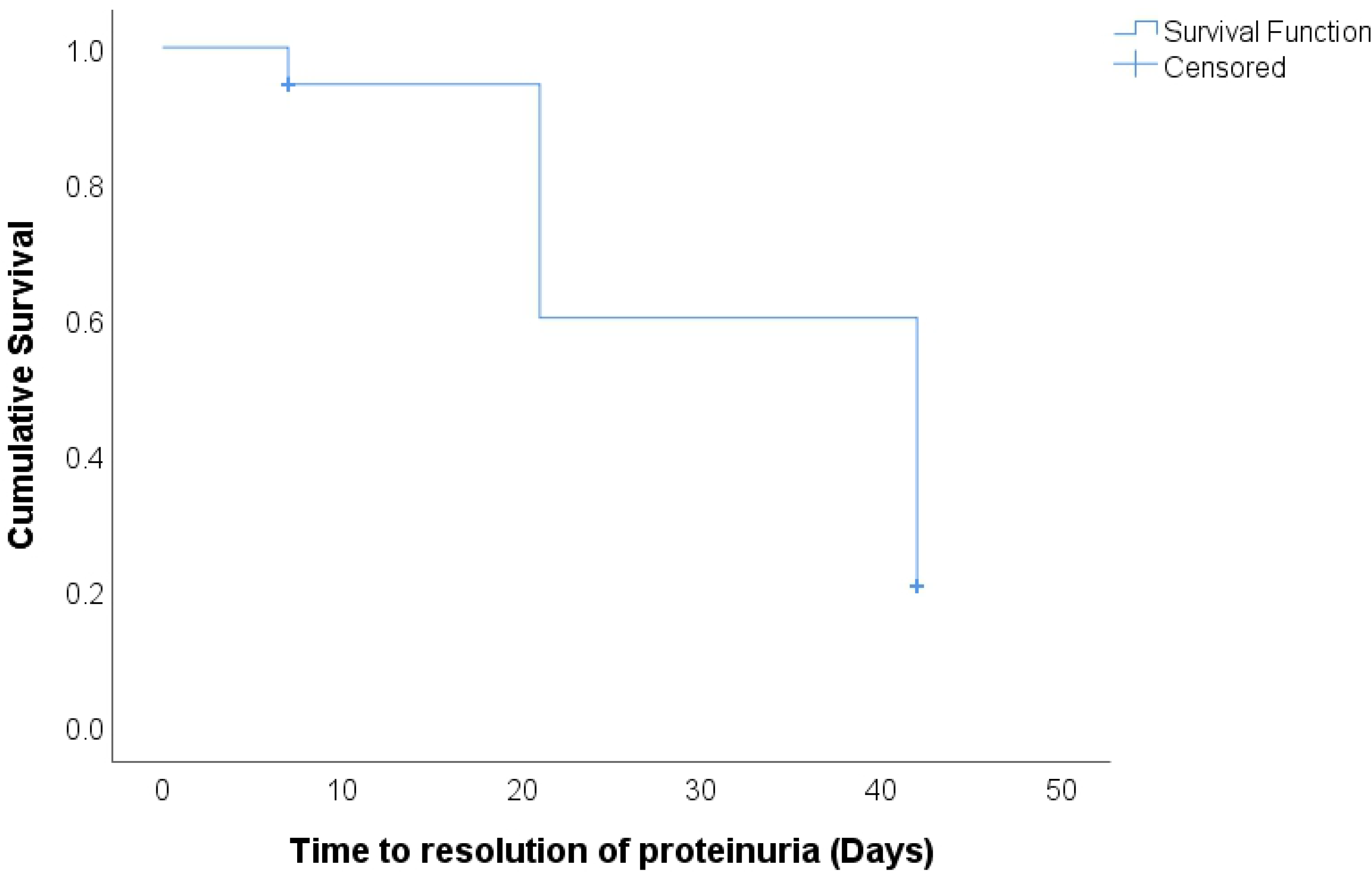
Kaplan-Meier curve of time to resolution of urine protein in women with severe preeclampsia and eclampsia.

Women with multiple pregnancy were more likely to have persistence of hypertension when compared women with singleton pregnancy (p=0.013)

## Discussion

This was a prospective cohort study in which 97 women with severe preeclampsia and eclampsia were followed after childbirth. In this study, 93.8% (91/97) of the women recovered from hypertension within 42 days after childbirth. This is in agreement with a study by Wei et al (18) in which 90% of their patients recovered from hypertension within 60 days after childbirth and Mikami et al (19) in which 90% of the women required 77 days to recover from hypertension. We considered the blood pressure to be normal if it was less than 140/90mmHg without using antihypertensive treatment for at least one week. In our study most women had an earlier recovery than the women in these other studies by Wei et al (18) and Mikami et al (19). This was probably because most mothers in our study had late onset preeclampsia at a gestational age of 35.9±4.0 weeks. Many studies have shown an association of a late onset of preeclampsia with early recovery of hypertension (11, 19). In our study 6 (6/97, 6.2%) women had persistent hypertension 6 weeks after delivery. This calls for prolonged follow up of women with hypertensive disease in pregnancy. Indeed the Society of Obstetricians and Gynaecologists of Canada clinical practice guidelines recommends follow up after six weeks to ascertain recovery from the effects of pregnancy and childbirth and ensure ongoing care with physicians or nephrologists(20). It has been shown that glomerular endothelial injury due to preeclampsia recovers within four weeks after delivery (21). Therefore women with persistent hypertension six weeks after delivery need adequate follow up to manage the underlying causes.

In this study, there was a statistically significant difference in the time to resolution of hypertension between singleton pregnancy versus multiple pregnancy: the time to resolution of hypertension was significantly shorter in singleton pregnancy (35.3±18.6 days) when compared to multiple pregnancy (43.5±31.4 days). This finding disagrees with what Mikami et. al. (19) found in their study. The normalization of blood pressure was significantly longer in singleton pregnancy than multiple pregnancy.

In this study, 47(47/97, 48.5%) participants had renal dysfunction after delivery: 24 women (24/47, 51.1%) were in stage 2, 18/47(38.3%) in stage 3 and 4/47(8.5%) in stage 4 of chronic kidney disease. At 6 weeks postpartum, 38 women (38/47, 80.9 %) resolved and 9(9/47, 19.1%) had persistent renal dysfunction: 6 in stage 2 and 3 in stage 3 chronic renal disease. The prevalence of renal dysfunction was higher than 24.1% that was found by Kaze et al in Cameroonian women (15) and 35.3% that was found by Prakash et al (6) in Indian women. Renal dysfunction is due glomerular endotheliosis which occurs in women with preeclampsia/eclampsia. Other studies have reported that resolution of the renal lesions may take up to two years (21). In our study, 18.6% of the women had persistent renal dysfunction six weeks after delivery. This is a reflection of the persistent effect of endothelial damage seen in preeclampsia(22). Most of these women are expected to recover within two years after delivery as this does not indicate chronic disease (23). However, this calls for prolonged follow up with a nephrologist.

We observed persistence of proteinuria of 16.3% at six weeks after delivery. This was lower than what was found in other studies (15, 19). Proteinuria is due endothelial dysfunction which plays a central role in the pathogenesis of preeclampsia (24). It has been shown that circulating soluble fms like tyrosine kinase-1 (sFlt-1) are elevated and these bind circulating vascular endothelial growth factor (VEGF). These have an association with glomerular endotheliosis and proteinuria(25). The resolution of proteinuria may echo the endothelial recovery seen after preeclampsia (26). These results may suggest that the time to resolution of proteinuria may be dependent on the degree and duration of endothelial cell injury. We expect most of the mothers with proteinuria to recover within two years as has been found in others studies(27). Proteinuria at six weeks does not indicate chronic disease. It most probably a temporary effect of endothelial damage seen in preeclampsia and more invasive investigations should be delayed until two years after delivery.

## Conclusion

In this study, 16.3% participants had persistent proteinuria, 6% had persistent hypertension and 19.6% had persistent renal function six weeks after delivery. Special interdisciplinary follow up of the patients with preeclampsia/eclampsia by an obstetrician and a physician after delivery is required to reduce maternal morbidity and mortality associated with preeclampsia/eclampsia in our community.

## Acknowledgements

Our thanks go to the staff of Mulago hospital maternal fetal medicine unit, the study participants, and our research team.

## Author Contributions

Conceived and designed the study: MK, PK, JB, JW, MWM. Participated in data collection and analysis: MK, PK, JB, JW, MWM, Interpreted the data and drafted the manuscript: MK, PK, JB, JW, MWM Reviewed and approved the final manuscript: MK, PK, JB, JW, MWM

